# Tracking the rates and mechanisms of canopy damage and recovery following Hurricane Maria using multitemporal lidar data

**DOI:** 10.1101/2021.03.26.436869

**Authors:** Veronika Leitold, Douglas C Morton, Sebastian Martinuzzi, Ian Paynter, Maria Uriarte, Michael Keller, António Ferraz, Bruce D Cook, Lawrence A Corp, Grizelle González

**Affiliations:** Department of Geographical Sciences, University of Maryland, College Park, 2181 LeFrak Hall, College Park, MD 20742 USA; Biospheric Sciences Laboratory, NASA Goddard Space Flight Center, 8800 Greenbelt Rd, Greenbelt, MD 20771 USA; SILVIS Lab, Department of Forest and Wildlife Ecology, University of Wisconsin-Madison, 1630 Linden Drive, Madison, WI 53706, USA; Universities Space Research Association, 7178 Columbia Gateway Dr, Columbia, MD 21046 USA; Department of Ecology, Evolution & Environmental Biology, Columbia University, 1190 Amsterdam Ave, New York, NY 10027; USDA-Forest Service International Institute of Tropical Forestry, Jardin Botanico Sur, 1201 Calle Ceiba, San Juan, PR 00926, USA; Jet Propulsion Laboratory, California Institute of Technology, 4800 Oak Grove Dr, Pasadena, CA 91109 USA; Institute of Environment and Sustainability, University of California, Los Angeles, LaKretz Hall, 619 Charles E Young Dr E #300, Los Angeles, CA 90024 USA; Science Systems and Applications Inc., 10210 Greenbelt Rd #600, Lanham, MD 20706 USA

**Keywords:** forest structure, cyclone, lidar, succession, ecosystem modeling, canopy traits, plasticity

## Abstract

Hurricane Maria (Category 4) snapped and uprooted canopy trees, removed large branches, and defoliated vegetation across Puerto Rico. The magnitude of forest damages and the rates and mechanisms of forest recovery following Maria provide important benchmarks for understanding the ecology of extreme events. We used airborne lidar data acquired before (2017) and after Maria (2018, 2020) to quantify landscape-scale changes in forest structure along a 439-ha elevational gradient (100 to 800 m) in the Luquillo Experimental Forest. Damages from Maria were widespread, with 73% of the study area losing ≥1 m in canopy height (mean = −7.1 m). Taller forests at lower elevations suffered more damage than shorter forests above 600 m. Yet only 13% of the study area had canopy heights ≤2 m in 2018, a typical threshold for forest gaps, highlighting the importance of damaged trees and advanced regeneration on post-storm forest structure. Heterogeneous patterns of regrowth and recruitment yielded shorter and more open forests by 2020. Nearly 45% of forests experienced initial height loss (<-1 m, 2017-2018) followed by rapid height gain (>1 m, 2018-2020), whereas 21.6% of forests with initial height losses showed little or no height gain, and 17.8% of forests exhibited no structural changes >|1| m in either period. Canopy layers <10 m accounted for most increases in canopy height and fractional cover between 2018-2020, with gains split evenly between height growth and lateral crown expansion by surviving individuals. These findings benchmark rates of gap formation, crown expansion, and canopy closure following hurricane damage.

**MANUSCRIPT HIGHLIGHTS:**

1. Hurricane Maria gave forests a haircut by toppling trees and shearing branches.
2. Regrowth after Maria was patchy, with equal areas of height gain and no change.
3. 3-D measures of forest recovery after hurricanes can improve ecosystem models.

## INTRODUCTION

Natural disturbances restructure forest ecosystems by altering the distributions of tree size, age, and species composition. In tropical and subtropical forests, cyclones are among the largest and most damaging disturbances. Cyclones occur with greater frequency than microburst “blow-down” events (Espírito-Santo and others, 2014) and cause widespread damage to coastal and inland tropical forests each year (e.g., Chen and others, 2015). Cyclonic storms are projected to increase in intensity (Knutson and others, 2010; Knutson and others, 2020) and frequency (Emanuel, 2013; Bhatia and others, 2018; but see Knutson and others, 2015) as a result of warming ocean temperatures (Hoyos and others, 2006) and rising atmospheric moisture content from climate change (Trenberth, 2018). Understanding the patterns and processes of forest disturbance and recovery following major cyclone events is therefore a priority to improve Earth system model predictions of the carbon, water, and energy balance of tropical forest regions (Seidl and others, 2011; U.S. DOE., 2018).

Hurricane Maria was the most powerful storm to hit Puerto Rico since 1928, making landfall just two weeks after Hurricane Irma on September 20, 2017. The Category 4 storm had maximum sustained winds of 210 km hr^-1^ (Van Beusekom and others, 2018), and the Luquillo Mountains in northeast Puerto Rico received >1200 mm of rainfall in just 48 hours (Hall and others, 2020).

Studies of initial forest damage at the individual-tree scale suggest that wind and rain from Hurricane Maria were more damaging to the forests of Puerto Rico than previous hurricanes (Tanner and others, 1991; Everham & Brokaw, 1996; Lugo, 2008; Uriarte and others, 2019). Hurricane damages to individual trees typically range from defoliation and branch fall to uprooting or stem breakage, with variability among tree species based on structural traits such as wood density, tree height, and crown size (Zimmerman and others, 1994; Canham and others, 2010). Hurricane Maria caused more stem breaks, even for species with high wood density, than Category 3 Hurricanes Hugo in 1989 or Georges in 1998 (Uriarte and others, 2019). The selective removal of taller individuals and canopy tree species with lower wood density favors palms and understory vegetation (Drew and others, 2009; Uriarte and others, 2019), leading to forests with lower canopies and higher stem densities (Ibanez and others, 2018). Delayed mortality of individuals damaged by Hurricane Maria, a process documented in past storms (Walker, 1995), could accentuate a long-term shift to fast-growing pioneer species and species most resilient to high winds, such as palms (Drew and others, 2009, Uriarte and others, 2021). Combined, Hurricanes Irma and Maria also caused a pulse of litter deposition that equaled or exceeded total annual litterfall, with a doubling of woody material (Liu and others, 2018).

Initial assessments of island-wide damages from Hurricane Maria using Landsat and Sentinel-2 satellite imagery data confirmed widespread defoliation. Patterns of defoliation manifested as a sharp decline in vegetation greenness, especially in the Luquillo Experimental Forest (Van Beusekom and others, 2018; Feng and others, 2020), and an increase in the sub-pixel fraction of non-photosynthetic vegetation, particularly in taller forests and areas that experienced high rainfall before and during Hurricane Maria (Hall and others, 2020). These studies also captured the heterogeneity of forest damages across Puerto Rico, consistent with the interactions between hurricane wind, rainfall, and local factors such as forest structure and topography (Tanner and others, 1991). However, passive optical satellite data are primarily sensitive to changes in fractional vegetation cover (Hu & Smith, 2018), and therefore do not differentiate defoliation from structural damages, or provide definitive evidence regarding the mechanisms of canopy recovery following hurricane disturbance (e.g., Feng and others, 2020).

Structural changes in the forest canopy from hurricane winds promote forest regeneration through recruitment, regrowth, or release (Everham & Brokaw, 1996; Uriarte and others, 2004; Uriarte and others, 2005; Drew and others, 2009; Shiels and others, 2015). Pioneer woody species can recruit from the seed bank immediately after the formation of a canopy opening, overtake existing non-pioneer trees based on rapid height growth, and dominate the adult community for many years following disturbances (Shiels and others, 2010). Smaller gaps in tropical forests typically lead to a combination of infilling and promotion from below, although the lateral growth of neighboring trees in tropical forests may be more limited (Hunter and others, 2015) than in temperate ecosystems (Runkle & Yetter, 1987; Young & Hubbell, 1991). In larger gaps, including in simulated hurricane experiments, defoliated or damaged stems often remain standing, limiting light penetration to the forest floor and potentially altering recovery pathways compared to non-hurricane gaps (Dietze & Clark, 2008). Damaged individuals can respond quickly after disturbance and flush new leaves or resprout within 7-10 weeks to rebuild the tree canopy, a process sometimes referred to as direct regeneration (Yih and others, 1991; Zimmerman and others, 1994; Tanner and others, 2014). Disturbance can also release surviving individuals in the canopy or understory from light, nutrient, or water competition, leading to rapid height growth (Uriarte and others, 2004; Shiels and others, 2010). Whether forest recovery proceeds via recruitment, regrowth, or release has distinct impacts on forest structure, and these different pathways affect the long-term changes in forest structure and function following hurricane damages.

Small-footprint airborne lidar data provide three-dimensional (3-D) information on forest structure needed to understand the underlying mechanisms that contribute to the structural reorganization of forests from hurricane damage and recovery. Multiple surveys of the same area capture fine-scale changes in canopy structure (e.g., Marvin & Asner, 2016; Leitold and others, 2018). Pre- and post-hurricane lidar data provide a unique canopy perspective on landscape-scale patterns. For example, the spatial and vertical distributions of residual canopy tree cover influence light availability and growing conditions for release and recruitment in the forest understory, and repeat measurements with high-density airborne lidar data capture height growth, crown expansion, and delayed treefall.

We analyzed a time series of airborne lidar data to quantify changes in forest structure from hurricane damages and recovery from Maria in the Luquillo Experimental Forest. Based on the elevational gradient (100-800 m) in pre-storm forest structure and species composition (Gould and others, 2006; Weaver, 2010) and post-storm research at the plot scale (Uriarte and others, 2019), we hypothesized that forest damage from Hurricane Maria would vary by forest type, with less damage in palm-dominated forests at higher elevations. However, the 3-D forest recovery following hurricane damages is poorly understood, based in part on competing processes of vegetation recovery and delayed mortality. Our two specific aims were therefore to 1) quantify landscape-scale variability in structural damage from Hurricane Maria, and 2) track the rates and mechanisms of changes in canopy structure during the first 2.5 years following the storm. Airborne data from NASA Goddard’s Lidar, Hyperspectral, and Thermal (G-LiHT) Airborne Imager were acquired over 439 ha, providing estimates of height changes across broad gradients of initial forest structure and composition. Understanding changes in forest structure from hurricane disturbance and forest recovery is necessary to capture the long-term effects of hurricanes on tropical forest ecosystems and improve Earth system models.

## METHODS

### Study area

The study area covers an elevational gradient on the northwest-facing slopes of the Luquillo Experimental Forest (coterminous with El Yunque National Forest) in northeast Puerto Rico (Figure 1). The climate is tropical maritime, with average annual rainfall of 3860 mm and average temperature of 22 °C in winter and 30 °C in summer (Quiñones and others, 2018). Elevation is the primary control on temperature and rainfall distributions, with differences of about 5 °C in mean annual temperature and >3000 mm in mean annual precipitation from the coast to the top of the Luquillo Mountains (González and others, 2013). The subtropical wet and rain forest formations within the study area are distributed by elevation in four main vegetation zones: (1) secondary wet forests in the lowlands; (2) tabonuco montane wet forests between 150-600 m dominated by tabonuco (*Dacryodes excelsa*) and motillo (*Sloanea berteriana*) trees with tall canopies reaching 30 m; (3) sierra palm (*Prestoea montana*) forests above 450 m that are common in steeper slopes and saturated soils; and (4) palo colorado (*Cyrilla racemiflora*) cloud forest between 600-900 m with tree heights reaching 15 m (Gould and others, 2006; Quiñones and others, 2018); our study area did not cover the elfin woodland vegetation type that is found on the tallest peaks above 900 m elevation. Hurricane Maria made landfall in Puerto Rico on 20 September 2017 as a Category 4 hurricane (Figure 1), damaging forests across the whole island, including the Luquillo Experimental Forest (Hall and others, 2020).

**Figure 1.**
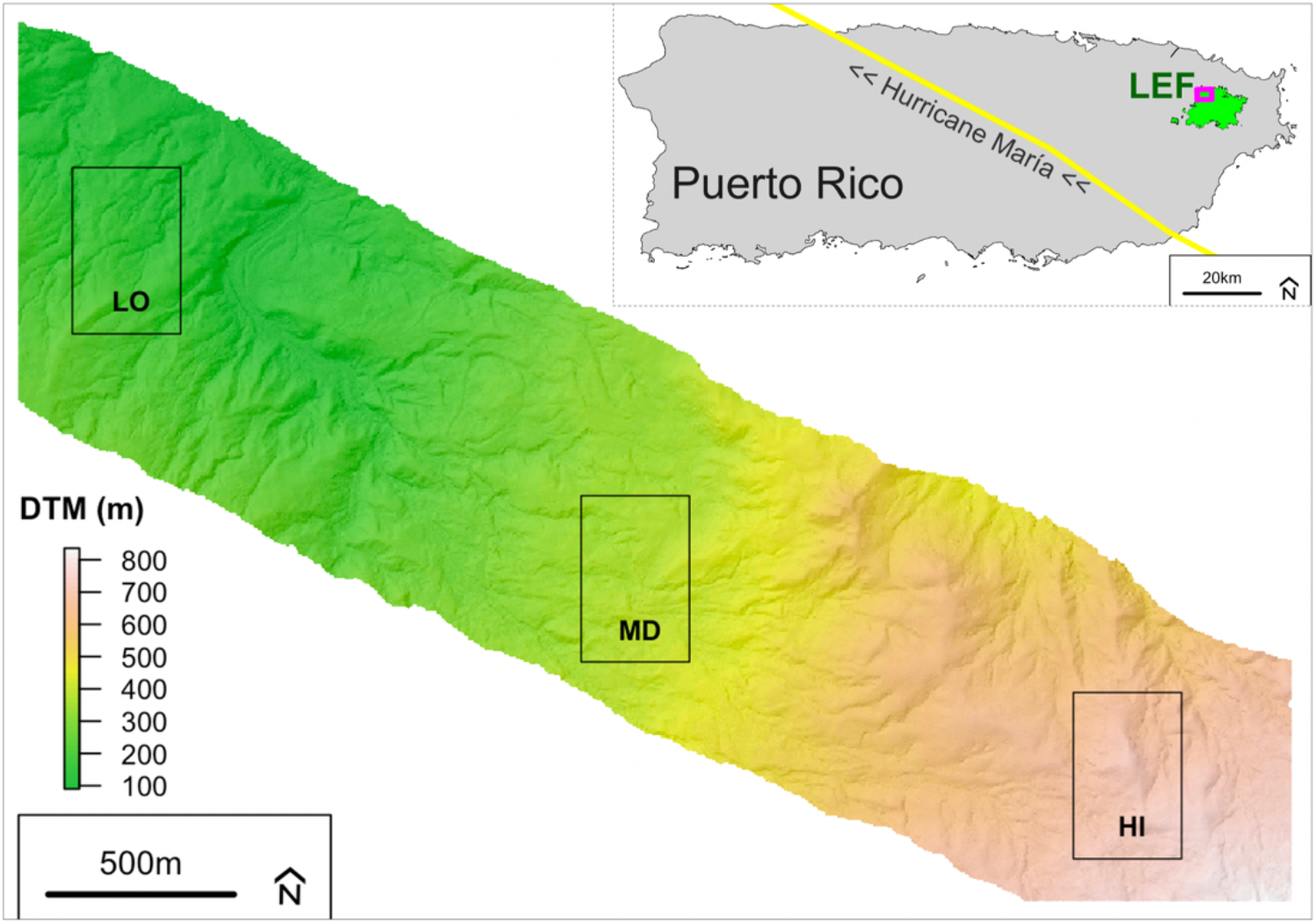
Digital terrain model (DTM) of the 439-ha study area showing the 100-800 m elevational gradient and the three 16-ha focus areas at low (LO), mid (MD), and high (HI) elevations. The inset panel (top right) shows the location of the study area (in magenta) within the Luquillo Experimental Forest (LEF) in northeast Puerto Rico and the path of Hurricane Maria (in yellow) across the island (NOAA, 2017).

### Airborne lidar data

Airborne lidar data over the Luquillo Experimental Forest were collected using the G-LiHT Airborne Imager (Cook and others, 2013) in three separate campaigns. Data were acquired in March 2017 (pre-hurricane), April 2018 (7 months post-hurricane), and March 2020 (2.5 years post-hurricane), with time intervals between lidar data collections of 13 months (2017-2018) and 23 months (2018-2020). All three airborne surveys used the same G-LiHT v2 instrumentation, including two VQ-480i scanning lidars (Riegl Laser Measurement Systems, Horn Austria). Data were acquired from a nominal flying altitude of 335 m AGL and 130 knots, which produced approximately 10 cm laser footprints (1550 nm) and an average sampling density of 12 laser pulses m^-2^. Pre- and post-flight boresight alignment of the lidar sensors ensured vertical accuracies of <10 cm (1 sigma) for all three campaigns. G-LiHT terrain and canopy height products are openly available online from the G-LiHT data portal (https://gliht.gsfc.nasa.gov).

The total area covered by all three lidar surveys was 439 ha, and subsets of the study area were designated using two different approaches (Figure 1). First, the study area was subdivided into 100-m terrain elevation classes to compare pre-hurricane forest structure, hurricane damages, and post-hurricane recovery along the gradient from 100 m to 800 m elevation. All G-LiHT DTM elevations were referenced to the EGM96 vertical datum. Second, we selected three focus areas at low (LO), mid (MD), and high (HI) elevations for analyses of changes in forest canopy structure over time, based on a consistent sample size. The mid-elevation (~380 m) focus area corresponds to the 16-ha Luquillo Forest Dynamics Plot (LFDP), part of the Long Term Ecological Research Network (LTER). This 16-ha rectangular area was replicated for the LO (~170 m elevation) and HI (~700 m elevation) focus areas, with the location of the LO and HI sites selected at random within areas with the lowest and highest elevations in the study area. The dominant vegetation type in the LO and MD focus areas is tabonuco forest, while the HI focus area contains palo colorado and sierra palm forests.

### Analysis

Lidar point clouds from the three airborne surveys (2017, 2018, and 2020) were processed using a consistent methodology (Cook and others, 2013) to evaluate changes in canopy height and canopy closure during the study period. Digital terrain models (DTM) and canopy height models (CHM) were generated separately for 2017, 2018 and 2020 at 1-m resolution. The decision to process lidar data from each campaign separately, rather than using a single reference DTM derived from all three campaigns, was based on the high lidar point density (Leitold and others, 2015) and the potential influence of landslides and erosion from the hurricane on the underlying topography (Van Beusekom and others, 2018; Bessette-Kirton and others, 2019). Changes in canopy height were calculated as the simple difference between CHM layers in 2017-2018, 2018-2020 and 2017-2020 at 1-m resolution. Canopy cover was estimated at each 1-m vertical increment above the ground as the percent ground area covered by vegetation at that height; cumulative canopy cover profiles were derived from these estimates. Similarly, canopy gap fraction was calculated at the native 1-m resolution of the lidar CHM layers as a measure of canopy openness—the percent ground area *not* covered by vegetation—at each 1-m vertical height bin in the canopy.

Forest canopy change was analyzed on a pixel-by-pixel basis and classified as one of three change categories: (a) *loss* pixels had canopy height change <-1 m between surveys, (b) *gain* pixels had canopy height change >1 m, and (c) *zero* pixels had canopy height change between −1 m and +1 m, i.e., a conservative estimate of near-zero changes in canopy height. Canopy height changes associated with hurricane damage (2017-2018) and post-hurricane recovery (2018-2020) created nine potential trajectories: loss-loss, loss-zero, loss-gain, zero-loss, zero-zero, zero-gain, gain-loss, gain-zero, and gain-gain. We summarized the magnitude of forest canopy changes in each trajectory on a per-area basis within the whole study area, in 100-m elevation bands (100-800 m), and in three 16-ha focus areas (LO, MD, HI). In each time interval, 1-m^2^ loss pixels were summed to estimate the fraction of forest area with canopy height losses. Previous studies have clustered loss pixels into individual canopy turnover events (Leitold and others, 2018). We therefore report annualized rates of canopy turnover using these more conservative thresholds for height loss (>3 m) and minimum size (>4 m^2^) for clusters of canopy turnover (Leitold and others, 2018). Change pixels were also grouped into vertical canopy height bins (1-m increments) and cohorts (5-m increments) to examine height changes and cumulative canopy cover within the forest profile during the study period. The relationship between pre-hurricane initial canopy height and post-hurricane height change was examined using ordinary least-square linear regression.

We analyzed patterns of vertical versus lateral (horizontal) growth in the canopy between 2018-2020 using a threshold for maximum vertical height gain during this interval derived from two approaches. First, we examined height gains within surviving canopy tree crowns in the three focus areas (LO, MD, HI). We used the ForestTools package (Plowright, 2020) of the R statistical software (R Core Team, 2020) to identify crown objects in the 2018 and 2020 canopy height models for trees at or exceeding the pre-storm mean canopy height. These canopy tree objects (containing one or more tree crowns) were used as inputs for the watershed segmentation algorithm to delineate the horizontal extent of all canopy tree crowns, not to separate individuals within the canopy stratum. We compared overlapping 2018 and 2020 canopy tree objects to estimate height gains within and adjacent to the 2018 crown extents (see Figure S1). Gains within the 2018 extent of crown objects were attributed to vertical growth, while height gains around crown edges were attributed to lateral expansion (Figure S1, Figure S2). Across the three focus areas, 85.1% - 89.3% of height gains within the extent of 2018 crown objects were ≤4 m.

Second, we compiled published values of the maximum height growth of pioneer tree species. We considered vertical growth of up to 4 m possible between 2018 and 2020, based on the maximum elongation of *Cecropia*, a dominant pioneer species in the understory (height growth up to 2.16 m yr^-1^ following disturbance, see Silander, 1979; also, Brokaw, 1998). Based on consistent estimates of vertical height growth for canopy trees and pioneer trees in the forest understory, we set a height gain threshold of 4 m to differentiate canopy closure from vertical growth (height gain ≤4 m) versus horizontal infilling by adjacent trees (height gain >4 m), and expressed these two different processes as percentages of the total area of canopy height gain.

## RESULTS

### Pre-Hurricane Forest Structure

In March 2017, closed-canopy tropical forests covered the entire study area (Figure 2).

**Figure 2.**
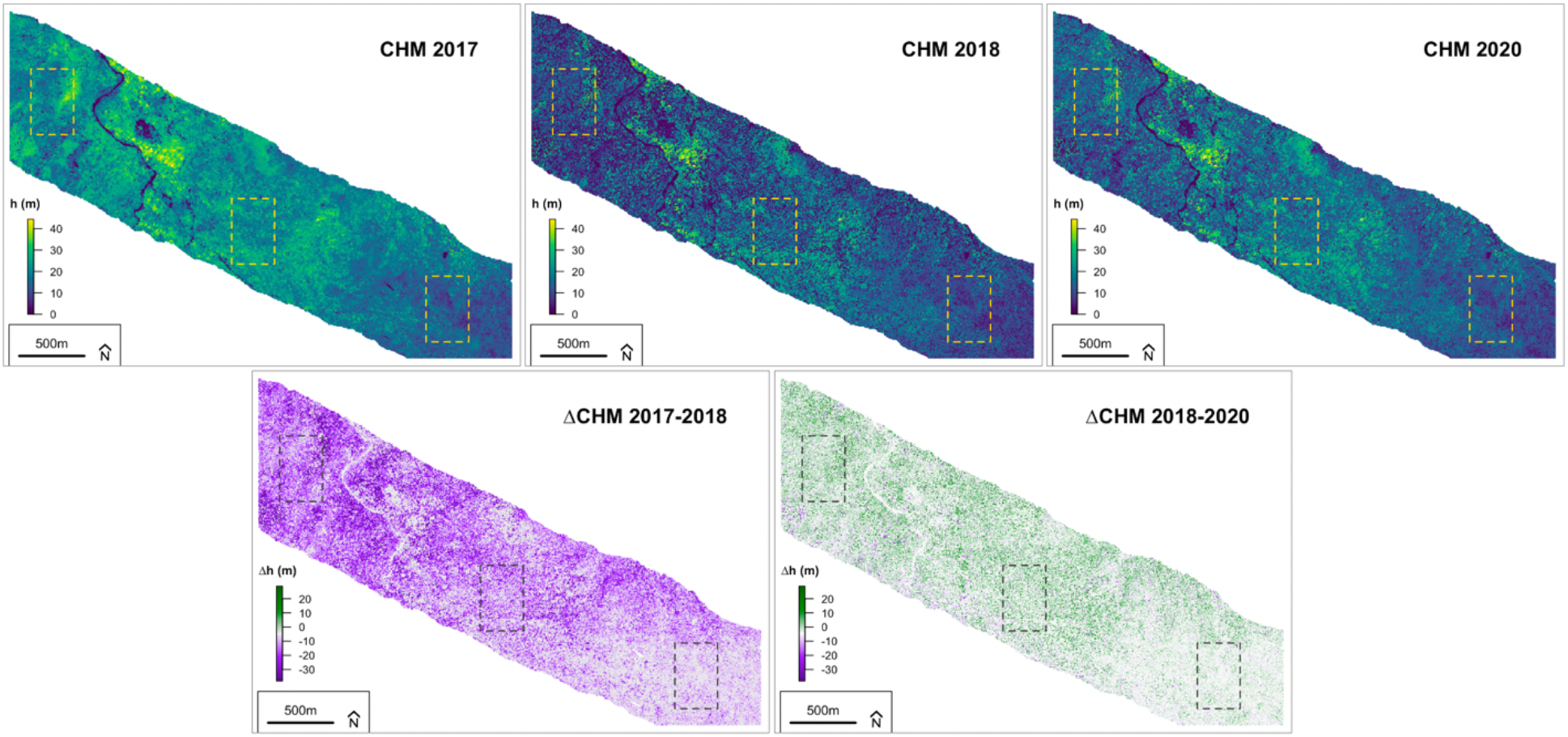
Canopy height models (CHM) at 1-m resolution of the study area in 2017, 2018, 2020 (top row), and canopy height change layers between 2017-2018 and 2018-2020 (bottom row). See supplemental Figure **S3** for the canopy height and changes in each 16-ha focus area.

The distribution of canopy heights was unimodal, with mean canopy height of 18.2±6.1 m (Table 1, Figure 3). Forests were tallest at lower elevations between 100-500 m, with mean canopy heights near 20 m. Forests above 600 m elevation had shorter trees with mean canopy heights of 13.8±3.8 m at 600-700 m and 10.9±3.4 m at 700-800 m. At all elevations, canopy cover was approximately 99% at 2 m and 98% at 5 m heights above the ground. For taller forests between 100-500 m, canopy cover was approximately 91% at 10 m above the ground.

**Table 1.**
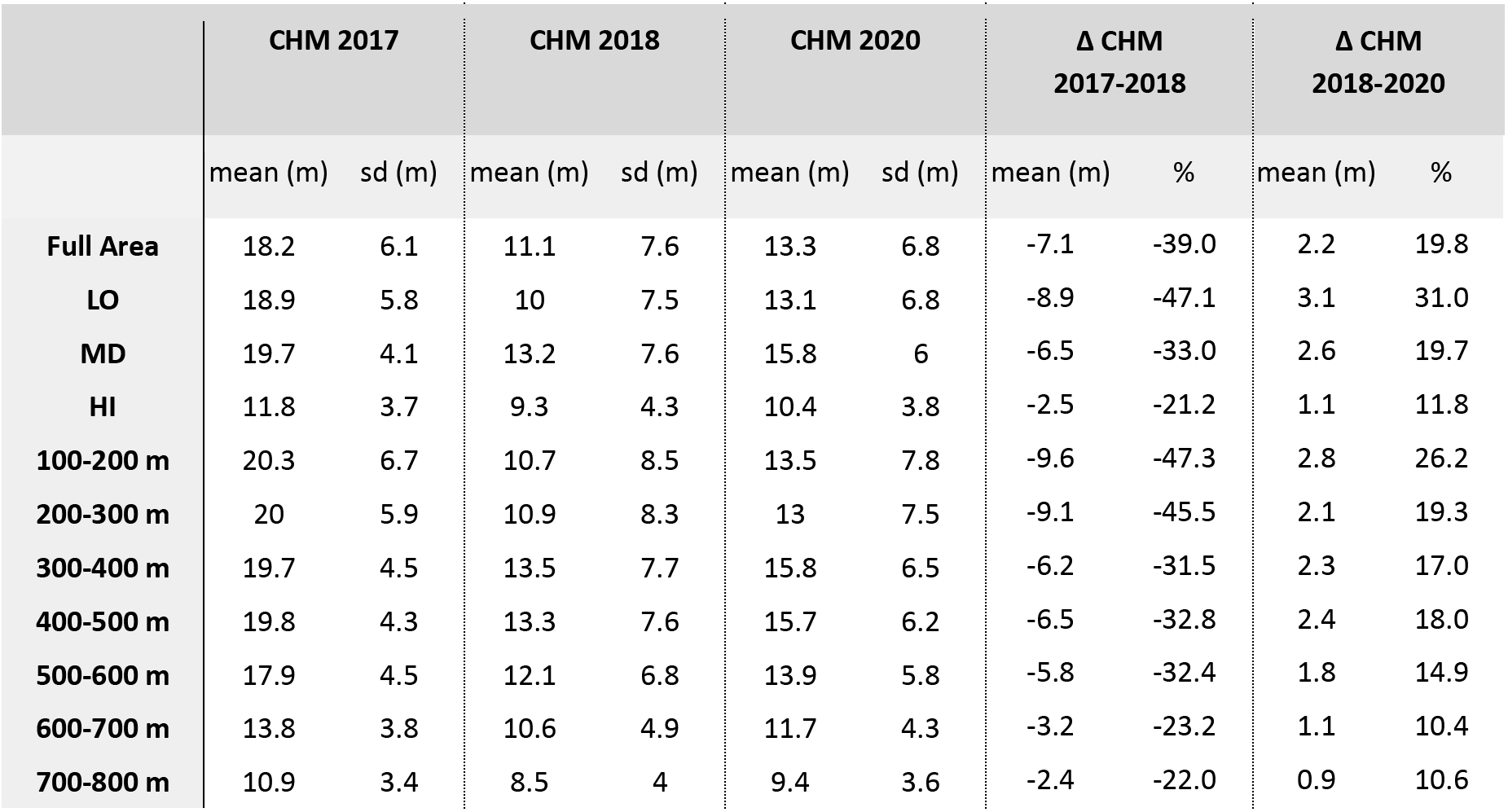
Canopy height and canopy height changes between 2017-2020 for the full study area, the 16-ha focus areas (LO, MD, HI), and the 100-m elevation bands. Mean and standard deviation (sd) canopy heights and canopy height changes were derived from the 2017, 2018, and 2020 Canopy Height Models (CHM) at 1-m spatial resolution. Focus area locations are shown in Figure 1.

**Figure 3.**
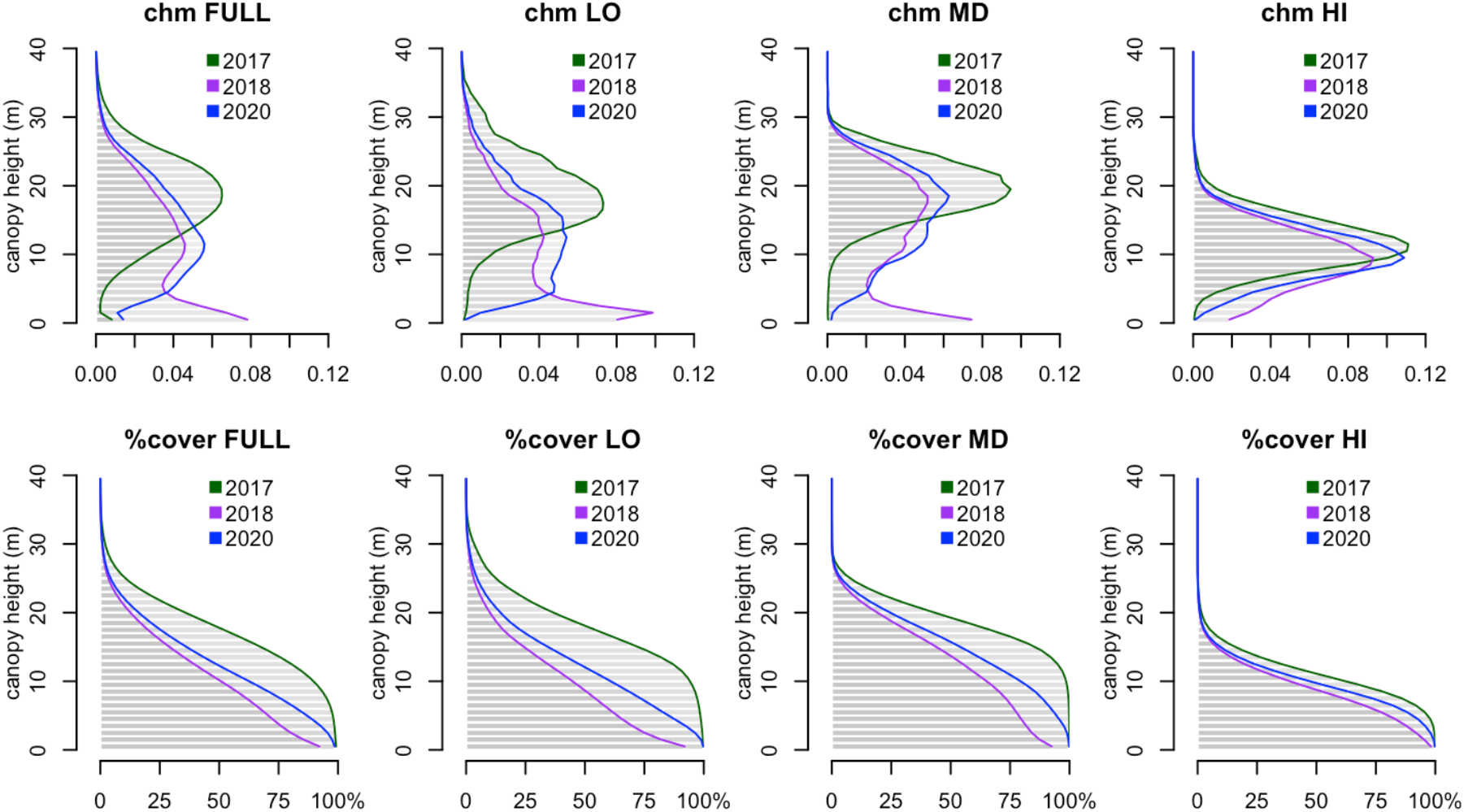
Vertical profiles of top of canopy height (top) and cumulative canopy cover (bottom) in 2017 (green), 2018 (magenta), and 2020 (blue) across the full study area (FULL) and in focus areas at low (LO), mid (MD) and high (HI) elevation.

### Hurricane Damage (2017-2018)

Forests were substantially shorter and more open seven months after Hurricane Maria (Table 1). Mean canopy height decreased by 7.1 m across the study region (39%), with average losses ranging from 5.8-9.6 m between 100-600 m elevation and 2.4-3.2 m above 600 m elevation. Mean height losses were strongly correlated with pre-storm mean canopy height (R^2^ = 0.80, p = 0.0063), as taller forests suffered larger height losses from Hurricane Maria. Declines in mean canopy height were largest at lower elevations, such that the tallest forests following the storm were at intermediate elevations (Figure S3). The distribution of canopy heights at low and middle elevations was bimodal in 2018, with one prominent mode corresponding to understory vegetation and advanced regeneration (1-2 m) and a second at the mean canopy height of 11.1±7.6 m (Figure 3).

Overall, approximately 73% of the study area lost at least 1 m of canopy height between the 2017 and 2018 lidar acquisitions (Table 2). Forests in the LO focus area had more damage, (80% ≥1 m canopy height loss), while only 54% of forests in the HI focus area had ≥1 m height loss from the storm (Figure S4). Based on a more conservative approach to cluster height losses into canopy turnover events using height-loss (>3 m) and size (>4 m^2^) thresholds from previous studies, canopy turnover from Hurricane Maria was 33 times higher than background canopy turnover in Amazon forests (1.8% yr^-1^, Leitold and others, 2018).

**Table 2.** Canopy change trajectories for the full study area and the three 16-ha focus areas (LO, MD, HI) expressed as percent area change in nine different change categories: loss-loss, loss-zero, loss-gain, zero-loss, zero-zero, zero-gain, gain-loss, gain-zero, and gain-gain. Gains and losses were separated from zero (no-change) areas based on >|1|m height changes.

|  |  | 2018 - 2020 |  |  |  | 2018 - 2020 |  |  |  | 2018 - 2020 |  |  |  | 2018 - 2020 |  |  |
| --- | --- | --- | --- | --- | --- | --- | --- | --- | --- | --- | --- | --- | --- | --- | --- | --- |
|  | Full Area | loss | zero | gain | LO | loss | zero | gain | MD | loss | zero | gain | HI | loss | zero | gain |
| 2017 - 2018 | loss | 6.37% | 21.59% | 44.62% | loss | 6.00% | 19.81% | 55.84% | loss | 4.90% | 22.09% | 46.41% | loss | 2.89% | 22.21% | 28.73% |
|  | zero | 2.86% | 17.82% | 3.25% | zero | 2.08% | 9.95% | 2.79% | zero | 2.65% | 18.81% | 2.86% | zero | 3.15% | 34.12% | 5.23% |
|  | gain | 0.95% | 2.06% | 0.43% | gain | 0.83% | 2.09% | 0.60% | gain | 0.60% | 1.52% | 0.27% | gain | 0.76% | 2.55% | 0.36% |

Height losses between 2017-2018 were concentrated in the dominant canopy layer around 20 m height, proportional to the distribution of pre-storm canopy heights (Figure 3). Branch loss and snapped or uprooted canopy trees therefore altered the vertical profile of canopy material, revealing layers of understory vegetation at lower canopy heights. The forest canopy in 2018 was significantly more open than before Maria, with 13.5% of top of canopy heights from 0-2 m above the ground, a typical height threshold for gaps in field studies (Brokaw, 1982), 25.7% ≤5 m, and 38.1% ≤10 m across the entire study area. Forests at 100-600 m elevation were significantly more open than forests at higher elevations, with approximately 15% and 5% canopy openings ≤2 m above the ground, respectively (Figure 3). Forests at lower elevations also had lower canopy cover throughout the vertical profile following the storm (e.g., 72% at 5 m, and 53% at 10 m).

### Post-Storm Recovery (2018-2020)

By March 2020, mean canopy height had increased from 11.1±7.6 m in 2018 to 13.3±6.8 m across the study area (Table 1). Average vertical height gains were greater at lower elevations (2-3 m) than above 600 m (~1 m), yet the tallest forests, on average, remained at intermediate elevations. Canopy closure increased rapidly between 2018 and 2020 at 2 m above the ground (97.5%) but open conditions persisted at 5 m (89%) and 10 m (65%) canopy heights (Figure 3). Approximately 50% of the landscape gained at least 1 m in canopy height during this 2.5-year interval (Table 2, Figure S4).

Between the 2018 and 2020 lidar acquisitions, approximately 10% of the study area lost >1 m height (5% yr^-1^). Clusters of canopy turnover based on the more conservative 3-m height loss and 4-m^2^ minimum size thresholds was 2.3% yr^-1^, more consistent with background turnover rates in other tropical forests (Leitold and others, 2018). Over 40% of the landscape had no detectable height change > |1 m| in the 2.5 years following the hurricane (Figure 4, Table 2), with canopy heights in these no-change areas concentrated around the pre-hurricane dominant tree height (mean heights of 16.8 m for the full study area, 18.8 m for LO, 20.7 m for MD, and 11.7 m for HI). Over the full study period (2017-2020), there was a net loss of canopy material at all canopy heights (Figure S4).

**Figure 4.**
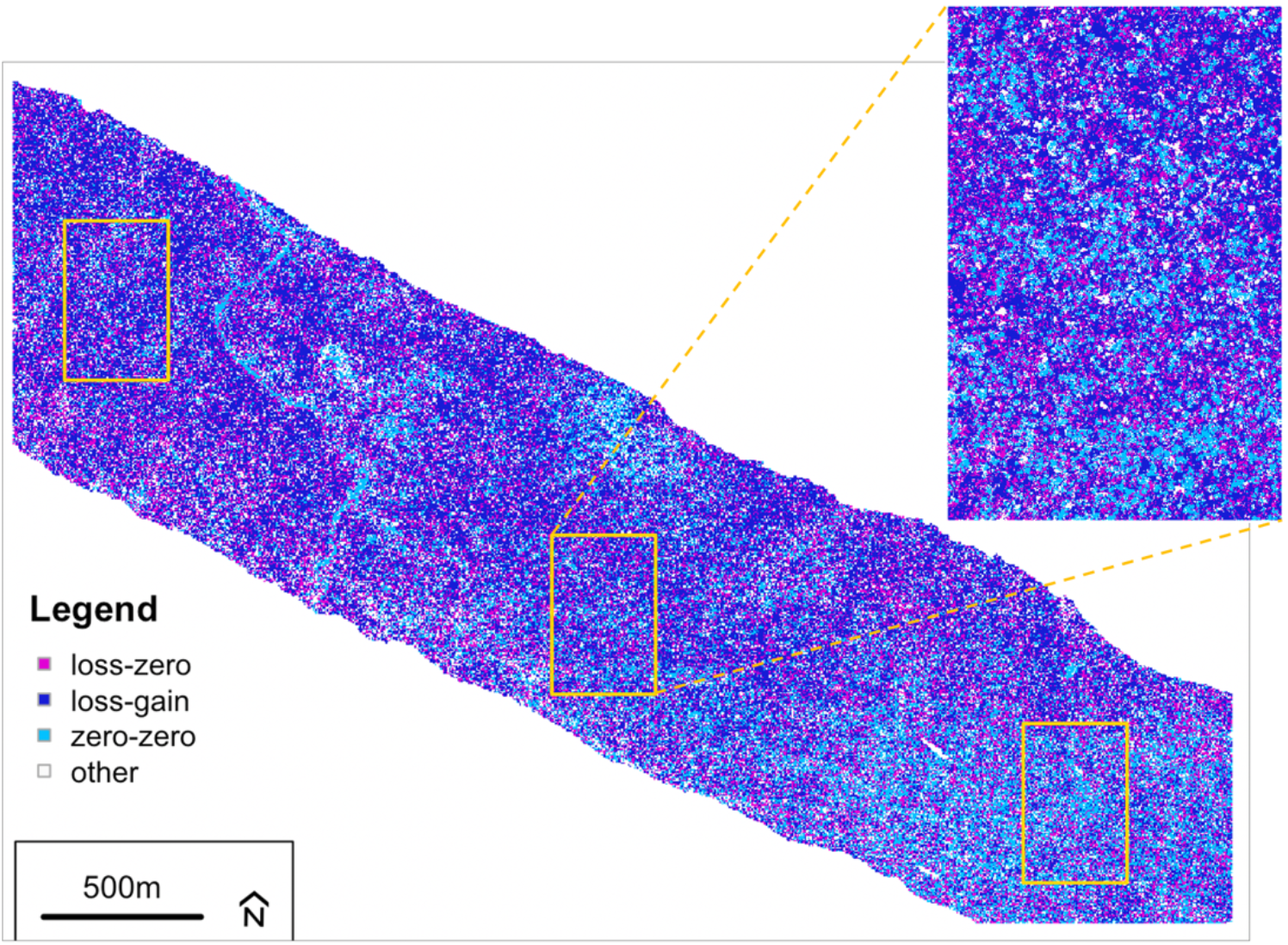
Map of the full study area showing the three dominant canopy change trajectories: loss-gain (44.6%), loss-zero (21.6%), and zero-zero (17.8%) at 1-m pixel resolution. All other change trajectories are shown in white (16%). The yellow rectangles indicate the locations of the three focus areas at low elevation (LO, top left), mid elevation (MD, center, and enlarged inset panel), and high elevation (HI, bottom right).

### Change trajectories

Three change trajectories of canopy damage and recovery were dominant at all elevations (Figure 4, Table 2). Nearly 45% of the study area followed the loss-gain trajectory, consistent with large height losses from the storm and rapid height gains between 2018 and 2020. The proportion of the loss-gain trajectory decreased with elevation from 56% (LO) to 29% (HI). The second most important trajectory was the loss-zero pathway (22%), suggesting delayed canopy recovery in areas of severe storm damage. The proportion of forest areas with loss-zero trajectories was similar across the elevation gradient in the full study area. The third most dominant trajectory was the zero-zero pathway (18%), forest canopy trees that neither lost height from the storm nor gained height from release in the post-storm environment. This trajectory was most common at high elevation (34%), where shorter trees, greater dominance of palms (Uriarte and others, 2019), and more regular exposure to strong winds may have resulted in both less storm damage and slower rates of post-storm height growth. Overall, the loss-gain and loss-zero trajectories were widely distributed across the study area, whereas the zero-zero trajectory exhibited clearer patterns at the individual tree crown and landscape scales (Figure 4).

The vertical distribution of canopy height changes provided important insight into the mechanisms of forest damage and recovery from Hurricane Maria (Figure 5). Across the entire study area, both height losses and zero (no-change) areas from 2017-2018 were proportional to the pre-storm distribution of canopy heights. In these loss areas, canopy material was concentrated below 5 m in height or distributed more uniformly between 5-30 m than in the pre-storm structure. Gains in canopy height between 2018 and 2020 created closed-canopy conditions at or below 5 m height and reestablished a dominant mode of canopy height at 12 m by 2020. Areas of no-change and loss between 2018 and 2020 occurred at all height levels, suggesting a highly heterogeneous canopy response.

**Figure 5.**
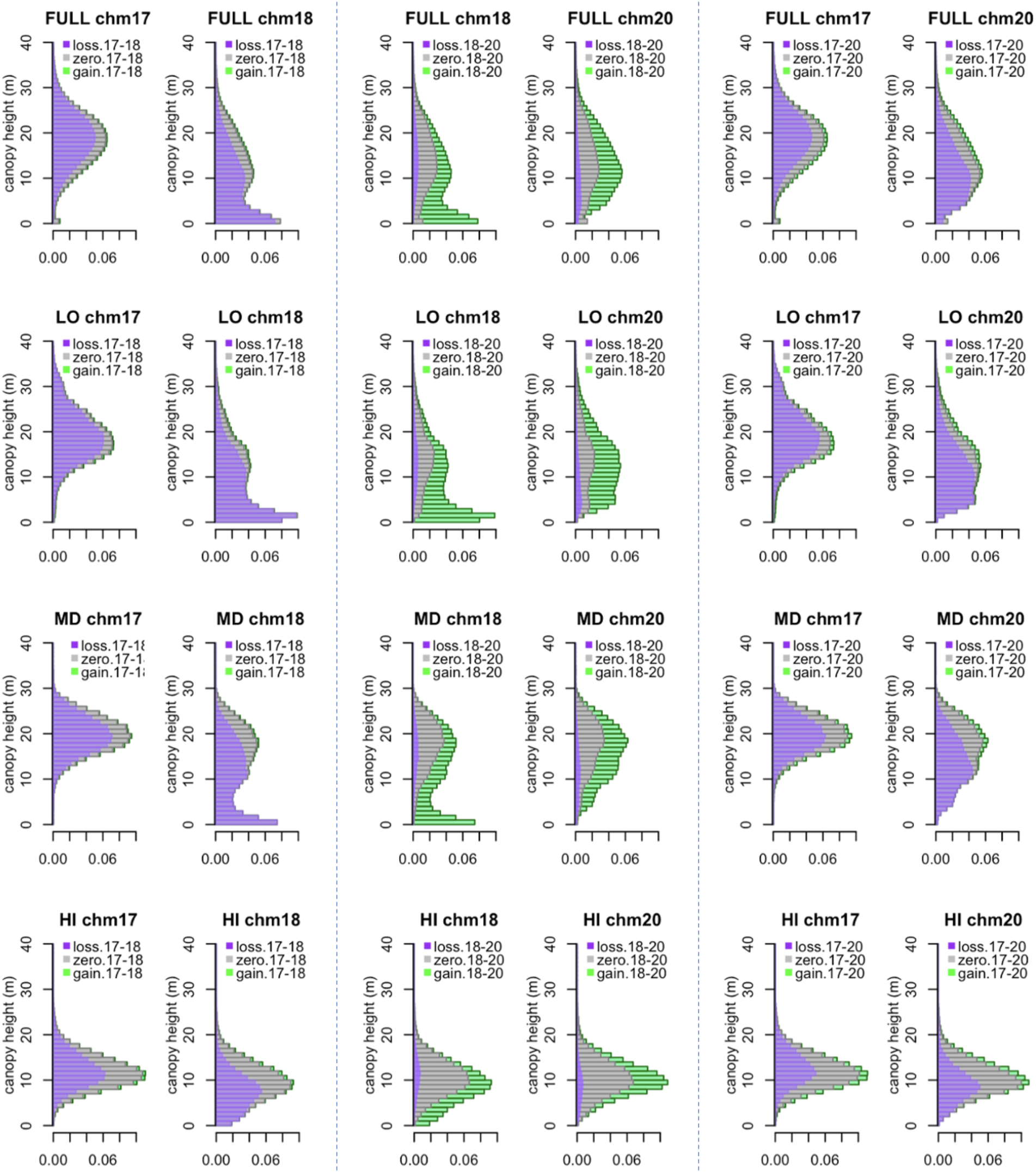
Vertical distribution of the changes in top of canopy heights, where paired figures in each time period illustrate the “from-to” movement of canopy material. Horizontal bars in each 1-m height bin are colored according to the proportion of area that falls within the height loss (purple), height gain (green), and zero change (gray) categories for each time interval.

The vertical redistribution of canopy material was similar in the LO and MD focus areas (Figure 5), and broadly consistent with the patterns observed across the entire study area. Nochange areas between 2017 and 2018 in the LO and MD focus areas tended to be in taller canopy positions, on average. These taller forest areas also suffered greater damages, with a bi-modal distribution of canopy heights in the loss category and the most open environments below 5 m. Shorter canopy environments closed rapidly between 2018-2020, with little or no replacement of gap areas with canopy height ≤2 m through further loss events.

Less damage at the HI focus area resulted in a more uniform vertical redistribution of canopy material. Canopy losses between 2017-2018 were proportional to the pre-storm canopy height distribution, but lower overall damages (Table 1) resulted in fewer canopy openings near the ground in 2018. Unlike the other focus areas, the most rapid growth response was near the mean canopy height. These factors limited overall changes in the vertical distribution of canopy material between 2017 and 2020 compared to portions of the study area at lower elevations.

Vertical height growth from established or new individuals and lateral growth of existing tree crowns contributed equally to canopy height gains between 2018 and 2020 for forest areas with loss-gain trajectories during the study period (Figure 5, Figure 6). Most height gains occurred in canopy positions <10 m in 2018. The shortest cohort (0-5 m) was more likely to be overtopped by neighboring trees (66%) than retain a canopy position through rapid height growth of new or established individuals (34%). For taller forests at LO and MD elevations, the canopy cohort from 5-10 m exhibited roughly equal proportions of height growth and infilling. For canopy positions >10 m in 2018, 73-88% of height gains were small, indicative of regrowth or expansion within existing crowns (≤2 m yr^-1^), with limited expansion of tree crowns by surviving individuals in the pre-storm canopy cohorts.

**Figure 6.**
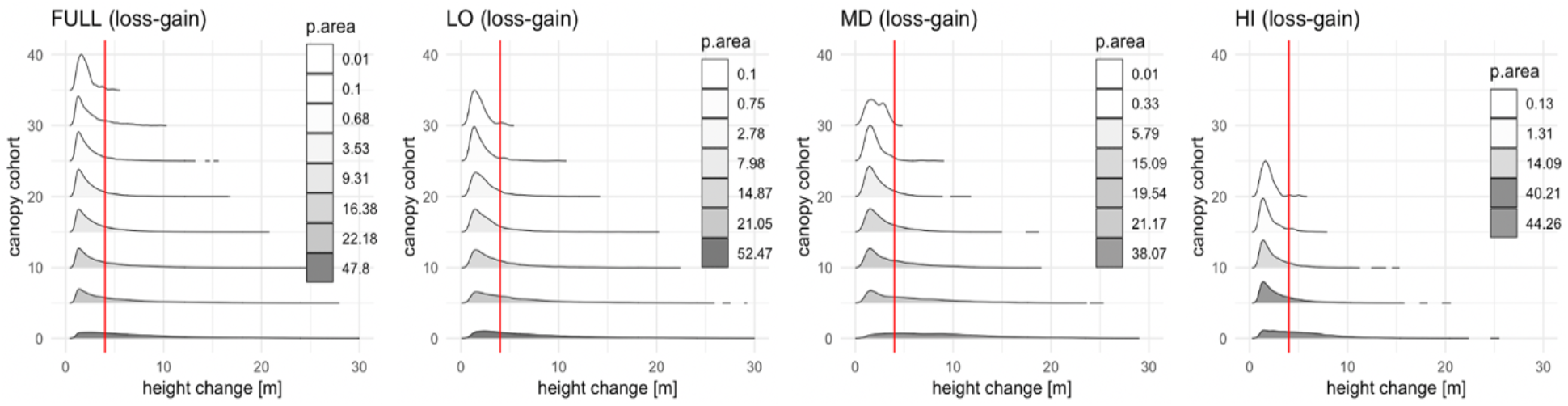
Distributions of canopy height gains between 2018 and 2020 for 5-m cohorts in forest areas with loss-gain trajectories in the full study area (FULL) and three 16-ha focus areas (LO, MD, HI). The vertical red lines indicate the threshold (4 m) separating vertical growth from horizontal infilling from neighboring trees, and shading denotes the proportion of all height gains in each 5-m cohort.

## DISCUSSION

Small footprint airborne lidar data captured widespread changes in forest structure from Hurricane Maria and rapid forest recovery along an elevational gradient of subtropical wet forest in Puerto Rico. Almost three-quarters of the study area lost canopy height from crown damage or treefall events. Yet, the pattern of hurricane damages was heterogeneous. Losses occurred at all canopy heights, and approximately one quarter of forests suffered little or no damage, including many taller and more exposed canopy positions. These findings reinforce the need to consider species-specific attributes (e.g., Uriarte and others, 2019) and topographic effects (Muscarella and others, 2020) in order to better understand the selective pressure of hurricanes on forest structure and composition. Nearly 2.5 years after the storm, forests were markedly shorter and more open, despite rapid vegetation growth in the forest understory. Surprisingly, more than 40% of forest canopy positions exhibited little or no height growth after the storm. Together, these nochange trajectories (loss-zero, zero-zero, gain-zero) were as abundant as forest areas with large height losses and subsequent height gains. Future work to combine lidar and inventory data may help reveal whether individuals with little or no height growth experienced lower productivity due to canopy damage, abiotic factors such as heat stress, or a shift in carbon allocation from height growth to other demands, such as fine root production (Silver & Vogt, 1993; Van Beusekom and others, 2020). Overall, these data provide important benchmarks of changes in forest structure from an extreme event, including critical inputs for ecosystem models regarding gap formation, initial and delayed structure changes, and the importance of surviving trees (with or without storm damage) on forest composition and productivity.

### Hurricane damage

Landscape-scale patterns of forest damage from repeat lidar data expand upon the understanding of hurricane damages from plot-scale studies. Brokaw & Grear (1991) estimated hurricane damage in 1-ha plots at three elevations in the Luquillo Mountains after Hurricane Hugo, documenting similar decreases in top of canopy height and variability in hurricane damages by forest type (21.1 m to 9.3 m in tabonuco and 10.1 m to 7.7 m in colorado forests). Lidar data in this study covered a much larger area (439 ha), capturing the spatial distributions of large and small height losses and the extent of forest patches with little or no damage from the hurricane. Lidar data revealed forest patches with lower hurricane damage (Figure 4), including well-drained sites at higher elevations. These findings clarify our understanding of fine-scale variability in hurricane damages across the landscape (Weaver, 2010) and motivate additional work on the complex interactions among rainfall, wind, and forest structure that create a mosaic of disturbance impacts from hurricanes.

Patterns of damage in this study differ from studies of initial forest damages from Hurricane Maria using moderate resolution satellite imagery (Van Beusekom and others, 2018; Feng and others, 2020; Hall and others, 2020). Lidar data provide precise, 3-D information at 1-m spatial resolution regarding the heterogeneity in canopy height loss, vertical distribution of vegetation within the forest profile, and fractional canopy cover. By contrast, estimates derived from optical satellite data estimate more uniform damages across the Luquillo Experimental Forest (Feng and others, 2020; Hall and others, 2020), despite general agreement for greater initial changes in NPV at low and mid elevations than for high-elevation forest types. Differences between lidar-derived changes in forest structure and optical measures of fractional vegetation cover suggest that initial defoliation is an imperfect proxy for structural damages from hurricane disturbance.

The spatial and vertical distributions of canopy openings following a major hurricane have not previously been quantified using lidar data. We estimated that Hurricane Maria generated canopy openings ≤2 m from branch loss and treefall events covering 13.5% of the study area, on average. Previous work at the plot scale reported even larger increases in gap area (vegetation height ≤2 m) from Hurricane Hugo in 1989 (Brokaw & Grear, 1991), with estimates of pre- and post-hurricane percent gap area of 0.4% versus 26% in tabonuco forest and 2% versus 27% in colorado forest, respectively. Results from this study for comparable forest types in the MD and HI focus areas had somewhat lower pre-hurricane gap area (0.05% and 0.14%) and substantially lower post-hurricane gap area (12.6% and 4.7% in tabonuco and colorado forests, respectively). Adding constraints for the minimum gap size, rather than summing all 1-m^2^ resolution pixels with canopy heights ≤2 m, would further reduce the estimate of gap formation from Hurricane Maria in our study area despite widespread evidence of crown damages and openings in the mid and upper canopy.

Three methodological factors may contribute to the large differences in estimated post-storm gap area between field and lidar studies. First, the time since hurricane may influence the estimates of gap area (9-20 weeks in Brokaw & Grear versus 7 months in this study), as resprouting and reflushing of canopy leaves may rapidly restore canopy cover (Tanner and others, 1991). Second, observer and laser-based estimates of forest structure differ substantially, even for common parameters such as tree height (e.g., Hunter and others, 2015). Brokaw & Grear (1991) collected field-based observations on a 5-m grid, with visual assessments of gap versus no-gap, compared to spatially and vertically-explicit measures of forest structure at 1-m resolution, with objective, laser-based estimates of canopy material. Third, estimates in this study reflect landscape-scale responses to hurricanes, not individual plots, given the extensive survey of 439 ha. However, direct comparisons between damage from different hurricanes are also confounded by other factions, including differences in pre-hurricane forest structure and composition, as selective removal of specific size and species classes from sequential storms (e.g., Uriarte and others, 2019) may reduce vulnerability at the stand scale in the short term.

One of the unique measurements from the time series of lidar data in this study is the direct comparison of initial and delayed canopy turnover following the storm. Previous studies have considered the long-term effects of other disturbance types using lidar data, including selective logging (Rangel Pinagé and others, 2019) and drought in Amazon forests (Leitold and others, 2018), but delayed structural damages from extreme events such as hurricanes have not been previously reported. We found limited evidence for delayed structural damages following the storm. While initial canopy turnover was 33 times higher than background turnover rates in Central Amazon forests (Leitold and others, 2018; Rangel Pinagé and others, 2019), canopy turnover between 2018-2020 in this study was comparable to background rates in these studies, and lower than annualized estimates of branch and treefall events in Western Amazon forests (Marvin & Asner, 2016).

At least three factors may contribute to the rapid return to background turnover following Hurricane Maria. First, structural damages within the first 7 months following the storm were included in the assessment of hurricane turnover, such that short-term delays in canopy turnover were attributed to initial forest damages. Second, canopy damages from Hurricane Maria reduced crown size for surviving individuals; the per-tree change in fractional canopy cover would therefore be smaller in the second time interval (2018-2020). Third, the survival rate of damaged individuals may be higher than expected, given the magnitude of initial canopy damages. Previous studies suggest that rates of tree mortality from hurricanes are low (<10% in Puerto Rican forests, see Frangi & Lugo 1991, Walker 1991, Zimmerman et al 1994, Uriarte and others 2019), even when entire forest stands suffer defoliation (Liu and others 2018) and large amounts of damage occur in the form of snapped or uprooted trees (Walker 1991, Basnet 1993, Vandermeer et al 1995). Overall, these findings suggest that hurricanes are a pulse disturbance, with direct impacts on individual tree and forest canopy structure, and long-term contributions from damaged canopy individuals to forest structure and function.

### Canopy recovery

Hurricane Maria created open-canopy conditions throughout the forest vertical profile, increasing light availability to spur rapid regrowth. Overall, nearly half of the forest area gained >1 m in height between 2018 and 2020. These height gains led to closure of canopy openings <10 m height from a combination of height growth of existing or new individuals and lateral crown expansion by surviving canopy trees or advanced regeneration. Repeat lidar measurements therefore suggest that rapid increases in leaf area captured by satellite vegetation indices (e.g., Feng and others, 2020) were concentrated in lower canopy layers, rather than from the rapid recovery of taller canopy trees.

We estimated that vertical and lateral growth contributed equally to canopy closure during this period, based on a threshold of maximum potential height growth (2 m yr^-1^), with marked differences in height growth distributions between taller and shorter cohorts. Importantly, the upper forest canopy remained open in 2020, with 65% canopy cover at 10 m compared to closed-canopy conditions at that height before the hurricane. Evidence for limited crown expansion from taller individuals (e.g., Figure 6) is consistent with previous reports of limited canopy tree plasticity adjacent to new gaps in other tropical forests (Hunter and others, 2015; Kellner & Asner, 2014). Rates of crown expansion may have been further suppressed by crown damage from Hurricane Maria, triggering epicormic resprouting from branch and stem positions lower in the forest vertical profile (Bellingham and others, 1994; Bellingham and others, 1996). Overall, this balance between vertical and horizontal growth has important implications for the size structure and species composition of hurricane-impacted forests (Uriarte and others, 2009; Uriarte and others, 2019). Our focus on canopy closure and increases in top of canopy height excluded important dynamics associated with recruitment and growth of understory vegetation that did not secure a canopy position following hurricane disturbance (Yih and others 1991, Vandermeer and others 1995, Uriarte and others, 2004; Dietze & Clark 2008; Shiels and others, 2010). Subsequent studies that consider the full lidar point cloud may offer insights regarding the spatial distribution and density of understory vegetation changes in response to initial changes in canopy structure from Hurricane Maria.

Even with high-density airborne lidar data, differentiating vertical growth from lateral expansion following Hurricane Maria remains a challenge. For example, we excluded subtle losses and gains of fine branches within tree crowns (<1 m height changes), potentially leading to an underestimate of height growth or release of surviving individuals. In addition, the use of a single threshold to separate height growth from crown expansion offers an initial estimate, whereas continuous distributions of height gains at all height levels illustrate the difficulty in separating specific mechanisms (e.g., Figure 6, Figure S2). Ideally, such a height threshold would be context-specific, based on observed differences in initial canopy height distributions and the presence or absence of advanced regeneration. Future studies that leverage other technologies may provide more confident separation of the mechanisms that contribute to structural changes in the post-hurricane environment. For example, terrestrial lidar scanners with higher point density and narrower beam divergence permit the separation of woody and leaf material, even for canopy vegetation in tall tropical forests (Boni Vicari and others, 2019). Similarly, volumetric analyses of tree objects (e.g., Ferraz and others, 2016; Ferraz and others, 2020), rather than the watershed approach used in this study, might also prove useful for evaluating the different processes of infilling within the original, pre-storm volumes occupied by canopy trees. Finally, field data also provide important constraints on the relative abundance of tree species that resprout following crown damage to guide the interpretation of lidar-derived estimates of changes in canopy structure.

### Key lessons for ecosystem models

Lidar-derived estimates of forest damage and recovery in this study provide four novel benchmarks for parameterization or validation of disturbance processes in ecosystem models. First, the current generation of ecosystem models represent the proportion of forest areas in different age classes since last disturbance (e.g., Fisher and others, 2018). Spatial and temporal variability in gap formation is therefore a critical parameter to ensure that long-term simulations capture the impact of hurricane disturbance. Here, we demonstrate that 13.5% of the study area had vegetation ≤2 m following the storm; model-dependent translation of this lidar-derived estimate would likely be lower, given minimum patch sizes in some models that build on the pseudo-spatial representation of forest structure first developed in the Ecosystem Demography model (e.g., Moorcroft and others, 2001; Fischer and others, 2016; Longo and others, 2020; Koven and others, 2020). Parameterization of new patches in size and age-structured models is an essential first step to improve the accuracy of post-disturbance recruitment following extreme events such as hurricanes. Second, trajectories of forest growth in ecosystem models typically vary based on plant functional types or plant traits, rather than disturbance. In this study, we estimate the proportion of the forested landscape that follows nine specific height gain and loss pathways from storm damage and initial forest recovery. These proportions, and the associated metrics of forest damage and recovery in each pathway, offer insights regarding damage-dependent rates and mechanisms of forest recovery, even for forests with similar species compositions and site conditions. Third, rates of canopy closure following hurricane disturbance at different heights within the forest vertical profile provide empirical values to constrain models that assume perfect plasticity of canopy material (e.g., Purves and others, 2008; Koven and others, 2020). Open canopy conditions in 2020 demonstrate that light penetration and growth of understory vegetation and shorter canopy cohorts constitute an important fraction of post-storm recovery. Initial findings of equal contributions from vertical and lateral growth also confirm the influence of light penetration for promotion of shorter cohorts. Finally, this study provides clear evidence for the long-term impact of damaged individuals on the structure and function of tropical forests following hurricane disturbance. Limited height growth, crown expansion, and turnover of surviving canopy trees during 2018-2020 underscore the longevity of damaged individuals in post-hurricane forest environments. Ultimately, it may be possible to use lidar-derived measures of forest structure to directly parameterize plant structural traits in ecosystem models to account for reductions in productivity and evapotranspiration for damaged individuals that differ from the standard allometric relationships in the model. Combined, these metrics of forest disturbance provide new constraints on forest growth and carbon cycling following hurricane damages, opening new avenues for future research.

## ACKNOWLEDGEMENTS

Funding for this study was provided by the US Department of Energy (Terrestrial Ecosystem Science Program, Interagency Agreements with the US Forest Service # 89243018SSC000012 and with NASA # 89243018SSC000013, and support to VL, DCM, and MK from the Next Generation Ecosystem Experiment-Tropics, Office of Biological and Environmental Research). Additional funding was provided by the USDA Forest Service, US Department of Interior (National Institute of Food and Agriculture # 2018-67030-28124), and NASA. The USDA Forest Service International Institute of Tropical Forestry, Luquillo LTER, and NASA’s Airborne Science Program provided logistical support.

## DATA AVAILABILITY

Lidar data in this study are online at https://gliht.gsfc.nasa.gov.

## AUTHOR CONTRIBUTIONS

VL and DCM designed the study; BDC and LAC collected airborne lidar data; VL and DCM analyzed data; all authors contributed to the interpretation of results and manuscript preparation.

## REFERENCES

Basnet K. 1993. Recovery of a Tropical Rain Forest after Hurricane Damage. Vegetation 109(1): 1–4.

Bellingham PJ, Tanner E, Healey J. 1994. Sprouting of Trees in Jamaican Montane Forests, after a Hurricane. Journal of Ecology 82(4): 747–758.

Bellingham PJ, Tanner EVJ, Rich PM, Goodland TCR. 1996. Changes in Light Below the Canopy of a Jamaican Montane Rainforest After a Hurricane. Journal of Tropical Ecology 12(5): 699–722.

Bessette-Kirton EK, Cerovski-Darriau C, Schulz WH, Coe JA, Kean JW, Godt JW, Thomas MA. 2019. Landslides triggered by hurricane Maria: An assessment of an extreme event in Puerto Rico. GSA Today, 29(6), 4–10.

Bhatia K, Vecchi G, Murakami H, Underwood S, Kossin J. 2018. Projected Response of Tropical Cyclone Intensity and Intensification in a Global Climate Model. Journal of Climate 31: 8281–8303.

Boni Vicari M, Disney MI, Wilkes P, Burt A, Calders K, Woodgate W. 2019. New framework for separating leaf and wood in terrestrial LiDAR point clouds. Methods in Ecology and Evolution 10: 680–694.

Brokaw NVL. 1998. Cecropia schreberiana in the Luquillo Mountains of Puerto Rico. Botanical Review 64: 91–120.

Brokaw NVL. 1982. The Definition of Treefall Gap and Its Effect on Measures of Forest Dynamics. Biotropica 14: 158–160.

Brokaw NVL, Grear J. 1991. Forest Structure Before and After Hurricane Hugo at Three Elevations in the Luquillo Mountains, Puerto Rico. Biotropica, 23(4): 386–392.

Canham CD, Thompson J, Zimmerman JK, Uriarte M. 2010. Variation in Susceptibility to Hurricane Damage as a Function of Storm Intensity in Puerto Rican Tree Species. Biotropica 42: 87–94.

Chen Y, Randerson JT, Morton DC. 2015. Tropical North Atlantic ocean-atmosphere interactions synchronize forest carbon losses from hurricanes and Amazon fires, Geophysical Research Letters 42: 6462–6470.

Cook BD, Corp LA, Nelson RF, Middleton EM, Morton DC, McCorkel JT, Masek JG, et al. 2013. NASA Goddard’s LiDAR, Hyperspectral and Thermal (G-LiHT) Airborne Imager. Remote Sensing 5(8): 4045–4066.

Dietze MC, Clark JS. 2008. Changing the gap dynamics paradigm: vegetative regeneration control on forest response to disturbance. Ecological Monographs 78: 331–347.

Drew AP, Boley JD, Zhao YH, Johnston MH, Wadsworth FH. 2009. Sixty-two years of change in subtropical wet forest structure and composition at El Verde, Puerto Rico. Interciencia 34: 34–40.

Emanuel KA. 2013. Downscaling CMIP5 climate models shows increased tropical cyclone activity over the 21st century, Proceedings of the National Academy of Sciences 110(30): 12219–12224.

Everham EM, Brokaw NVL. 1996. Forest Damage and Recovery from Catastrophic Wind. The Botanical Review 62: 113–185.

Espírito-Santo FDB, Gloor M, Keller M, Malhi Y, Saatchi S, Nelson B, Oliveira Junior RC, Pereira C, Lloyd J, Frolking S, Palace M, Shimabukuro YE, Duarte V, Mendoza AM, López-González G, Baker TR, Feldpausch TR, Brienen RJW, Asner GP, Boyd DS, Phillips OL. 2014. Size and frequency of natural forest disturbances and the Amazon forest carbon balance. Nature Communications 5:3434.

Feng Y, Negrón-Juárez R I, Chambers JQ. 2020. Remote sensing and statistical analysis of the effects of hurricane María on the forests of Puerto Rico, Remote Sensing of Environment 247: 111940.

Ferraz A, Saatchi S, Mallet C, Meyer V. 2016. Lidar detection of individual tree size in tropical forests. Remote Sensing of Environment 183: 318–333.

Ferraz A, Saatchi SS, Longo M, Clark DB. 2020. Tropical tree size-frequency distributions from airborne lidar. Ecological Applications 30(7): e02154.

Fischer R, Bohn F, de Paula MD, Dislich C, Groeneveld J, Gutiérrez AG, Kazmierczak M, Knapp N, Lehmann S, Paulick S, Pütz S, Rödig E, Taubert F, Köhler P, Huth A. 2016. Lessons learned from applying a forest gap model to understand ecosystem and carbon dynamics of complex tropical forests. Ecological Modelling, 326: 124–133.

Fisher RA, Koven CD, Anderegg WRL, Christoffersen BO, Dietze MC, Farrior CE, Holm JA, Hurtt GC, Knox RG, Lawrence PJ, Lichstein JW, Longo M, Matheny AM, Medvigy D, Muller-Landau HC, Powell TL, Serbin SP, Sato H, Shuman JK, Smith B, Trugman AT, Viskari T, Verbeeck H, Weng E, Xu C, Xu X, Zhang T, Moorcroft PR. 2018. Vegetation demographics in Earth System Models: A review of progress and priorities. Global Change Biology 24: 35–54.

Frangi J, Lugo A. 1991. Hurricane Damage to a Flood Plain Forest in the Luquillo Mountains of Puerto Rico. Biotropica 23(4): 324–335.

González G, Waide RB, Willig MR. 2013. Advancements in the understanding of spatiotemporal gradients in tropical landscapes: a Luquillo focus and global perspective. Ecological Bulletins 54: 245–250.

Gould WA, González G, Carrero Rivera G. 2006. Structure and composition of vegetation along an elevational gradient in Puerto Rico. Journal of Vegetation Science 17: 563–574

Hall J, Muscarella R, Quebbeman A, Arellano G, Thompson J, Zimmerman JK, Uriarte M. 2020. Hurricane-Induced Rainfall is a Stronger Predictor of Tropical Forest Damage in Puerto Rico Than Maximum Wind Speeds. Scientific Reports 10: 4318.

Hoyos CD, Agudelo PA, Webster PJ, Curry JA. 2006. Deconvolution of the factors contributing to the increase in global hurricane intensity. Science 312: 94–97.

Hu T, Smith RB. 2018. The Impact of Hurricane Maria on the Vegetation of Dominica and Puerto Rico Using Multispectral Remote Sensing. Remote Sensing 10: 827.

Hunter MO, Keller M, Morton D, Cook B, Lefsky M, Ducey M, Saleska S, Oliveira RC, Schietti J. 2015. Structural Dynamics of Tropical Moist Forest Gaps. PLoS ONE 10(7): 0132144.

Kellner JR, Asner GP. 2014. Winners and losers in the competition for space in tropical forest canopies. Ecology Letters 17(5): 556–562.

Knutson T, McBride JL, Chan J, Emanuel K, Holland G, Landsea C, Held I, Kossin JP, Srivastava AK, Sugi M. 2010. Tropical cyclones and climate change. Nature Geoscience 3: 157–163.

Knutson TR, Sirutis JJ, Zhao M, Tuleya RE, Bender M, Vecchi GA, Villarini G, Chavas D. 2015. Global Projections of Intense Tropical Cyclone Activity for the Late Twenty-First Century from Dynamical Downscaling of CMIP5/RCP4.5 Scenarios. Journal of Climate 28: 7203–7224.

Knutson T, Camargo SJ, Chan JCL, Emanuel K, Ho CH, Kossin J, Mohapatra M, Satoh M, Sugi M, Walsh K, Wu L. 2020. Tropical Cyclones and Climate Change Assessment: Part II: Projected Response to Anthropogenic Warming. Bulletin of the American Meteorological Society 101: E303–E322.

Koven CD, Knox RG, Fisher RA, Chambers JQ, Christoffersen BO, Davies SJ, Detto M, Dietze MC, Faybishenko B, Holm J, Huang M, Kovenock M, Kueppers LM, Lemieux G, Massoud E, McDowell NG, Mueller-Landau HC, Needham JF, Norby RJ, Powell T, Rogers A, Serbin SP, Shuman JK, Swann AL, Varadharajan C, Walker AP, Wright SJ, Xu C. 2020. Benchmarking and parameter sensitivity of physiological and vegetation dynamics using the Functionally Assembled Terrestrial Ecosystem Simulator (FATES) at Barro Colorado Island, Panama. Biogeosciences 17: 3017–3044.

Leitold V, Keller M, Morton DC, Cook BD, Shimabukuro YE. 2015. Airborne lidar-based estimates of tropical forest structure in complex terrain: opportunities and trade-offs for REDD+. Carbon Balance and Management 10(1): 3.

Leitold V, Morton DC, Longo M, dos-Santos MN, Keller M, Scaranello M. 2018. El Niño drought increased canopy turnover in Amazon forests. New Phytologist 219(3): 959–971.

Liu X, Zeng X, Zou X, González G, Wang C. Yang S. 2018. Litterfall Production Prior to and during Hurricanes Irma and Maria in Four Puerto Rican Forests. Forests 9(6): 367–.

Longo M, Saatchi S, Keller M, Bowman K, Ferraz A, Moorcroft PR, Morton DC, Bonal D, Brando P, Burban B, Derroire G, dos-Santos MN, Meyer V, Saleska S, Trumbore S, Vincent G. 2020. Impacts of degradation on water, energy, and carbon cycling of the Amazon tropical forests. Journal of Geophysical Research Biogeosciences 125: e2020JG005677.

Lugo AE. 2008. Visible and invisible effects of hurricanes on forest ecosystems: an international review. Austral Ecology 33: 368–398.

Marvin DC, Asner GP. 2016. Branchfall dominates annual carbon flux across lowland Amazonian forests. Environmental Research Letters 11: 094027.

Moorcroft PR, Hurtt GC, Pacala SW. 2001. A method for scaling vegetation dynamics: The Ecosystem Demography model (ED). Ecological Monographs, 71(4), 557–586.

Muscarella R, Kolyaie S, Morton DC, Zimmerman JK, Uriarte M. 2020. Effects of topography on tropical forest structure depend on climate context. Journal of Ecology 108: 145–159.

National Oceanic and Atmospheric Administration (NOAA). 2017. National Hurricane Center, Tropical Cyclone Report: Hurricane Maria. Available online at: https://www.nhc.noaa.gov/data/tcr/AL152017_Maria.pdf. Accessed November 13, 2020.

Plowright A. 2020. ForestTools: Analyzing Remotely Sensed Forest Data. R package version 0.2.1. Available online at: https://cran.r-project.org/web/packages/ForestTools/ForestTools.pdf. Accessed November 13, 2020.

Purves DW, Lichstein JW, Strigul N, Pacala SW. 2008. Predicting and understanding forest dynamics using a simple tractable model. Proceedings of the National Academy of Sciences 105(44): 17018–17022.

Quiñones M, Parés-Ramos IK, Gould WA, González G. McGinley K, Ríos P. 2018. El Yunque National Forest Atlas. General Technical Report IITF-GTR-47. San Juan, PR: U.S. Department of Agriculture, Forest Service, International Institute of Tropical Forestry. 63p.

R Core Team. 2020. R: A language and environment for statistical computing. v.3.6.2. R Foundation for Statistical Computing, Vienna, Austria. URL http://www.R-project.org/.

Rangel Pinagé E, Keller M, Duffy P, Longo M, dos-Santos MN, Morton DC. 2019. Long-Term Impacts of Selective Logging on Amazon Forest Dynamics from Multi-Temporal Airborne LiDAR. Remote Sensing 11(6): 709

Runkle J, Yetter T. 1987. Treefalls Revisited: Gap Dynamics in the Southern Appalachians. Ecology 68(2): 417–424.

Seidl R, Fernandes PM, Fonseca TF, Gillet F, Jonsson AM, Merganicova K, Netherer S, Arpaci A, Bontemps JD, Bugmann H, Gonzalez-Olabarria JR, Lasch P, Meredieu C, Moreira F, Schelhaas MJ, Mohren F. 2011. Modelling natural disturbances in forest ecosystems: a review. Ecological Modelling 222: 903–924.

Shiels AB, Zimmerman JK, García-Montiel DC, Jonckheere I, Holm J, Horton D, Brokaw N. 2010. Plant responses to simulated hurricane impacts in a subtropical wet forest, Puerto Rico. Journal of Ecology 98(3): 659–673.

Shiels AB, González G, Lodge DJ, Willig MR, Zimmerman JK. 2015. Cascading Effects of Canopy Opening and Debris Deposition from a Large-Scale Hurricane Experiment in a Tropical Rain Forest. BioScience 65(9): 871–881.

Silander SR. 1979. A study of the ecological life history of Cecropia peltata L., an early secondary successional species in the rain forest of Puerto Rico. Thesis (M.S.), University of Tennessee, Institute of Ecology, Knoxville. 94p.

Silver WL, Vogt KA. 1993. Fine Root Dynamics Following Single and Multiple Disturbances in a Subtropical Wet Forest Ecosystem. Journal of Ecology 81(4): 729–738.

Tanner EVJ, Kapos V, Healey JR. Hurricane Effects on Forest Ecosystems in the Caribbean. 1991. Biotropica 23(4): 513–21.

Tanner EVJ, Rodriguez-Sanchez F, Healey JR, Holdaway RJ, Bellingham PJ. 2014. Long-term hurricane damage effects on tropical forest tree growth and mortality. Ecology 95: 2974–2983.

Trenberth KE, Cheng L, Jacobs P, Zhang Y, Fasullo J. 2018. Hurricane Harvey links to ocean heat content and climate change adaptation. Earth’s Future, 6: 730–744.

Uriarte M, Canham CD, Thompson J, Zimmerman JK. 2004. A Neighborhood analysis of tree growth and survival in a hurricane-driven tropical forest. Ecological Monographs 74: 591–614.

Uriarte M, Canham CD, Thompson J, Zimmerman JK, Brokaw N. 2005. Seedling recruitment in a hurricane driven tropical forest: light limitation, density-dependence and the spatial distribution of parent trees. Journal of Ecology 93: 291–304.

Uriarte M, Canham CD, Thompson J, Zimmerman JK, Murphy L, Sabat AM, Fetcher N, Haines BL. 2009. Natural disturbance and human land use as determinants of tropical forest dynamics: results from a forest simulator. Ecological Monographs 79: 423–443.

Uriarte M, Thompson J, Zimmerman JK. 2019. Hurricane Mariia tripled stem breaks and doubled tree mortality relative to other major storms. Nature Communications 10: 1362.

Uriarte M, Tang C, Morton DC, Zimmerman JK, Zheng T. 2021. Hurricanes leave long lasting legacies on tropical forest structure and composition. Nature Ecology & Evolution, submitted.

U.S. Department of Energy (U.S. DOE). 2018. Disturbance and Vegetation Dynamics in Earth System Models; Workshop Report, DOE/SC-0196. Office of Biological and Environmental Research, U.S. Department of Energy Office of Science. Available online at https://tes.science.energy.gov/files/vegetationdynamics.pdf. Accessed November 13, 2020.

Van Beusekom AE, Álvarez-Berríos NL, Gould WA, Quiñones M, González G. 2018. Hurricane Maria in the U.S. Caribbean: Disturbance forces, variation of effects, and implications for future storms. Remote Sensing 10: 1–14.

Van Beusekom AE, González G, Stankavich S, Zimmerman JK, Ramírez A. 2020. Understanding tropical forest abiotic response to hurricanes using experimental manipulations, field observations, and satellite data. Biogeosciences 17(12): 3149–3163.

Vandermeer J, Mallona M, Boucher D, Yih K, Perfectos I. 1995. Three Years of Ingrowth Following Catastrophic Hurricane Damage on the Caribbean Coast of Nicaragua: Evidence in Support of the Direct Regeneration Hypothesis. Journal of Tropical Ecology 11(3): 465–471.

Walker L. 1991. Tree Damage and Recovery from Hurricane Hugo in Luquillo Experimental Forest, Puerto Rico. Biotropica 23(4): 379–385.

Walker LR. 1995. Timing of post-hurricane tree mortality in Puerto Rico. Journal of Tropical Ecology 11: 315–320.

Weaver PL. 2010. Forest Structure and Composition in the Lower Montane Rain Forest of the Luquillo Mountains, Puerto Rico. Interciencia 35: 640–646.

Yih K, Boucher D, Vandermeer J, Zamora N. 1991. Recovery of the Rain Forest of Southeastern Nicaragua After Destruction by Hurricane Joan. Biotropica 23(2): 106–113.

Young T, Hubbell S. 1991. Crown Asymmetry, Treefalls, and Repeat Disturbance of Broad-Leaved Forest Gaps. Ecology 72(4): 1464–1471.

Zimmerman JK, Everham EM, Waide RB, Lodge DJ, Taylor CM, Brokaw NVL. 1994. Responses of Tree Species to Hurricane Winds in Subtropical Wet Forest in Puerto-Rico: Implications for Tropical Tree Life Histories. Journal of Ecology 82: 911–922.

