## Supplemental Figures for "Tracking the rates and mechanisms of canopy damage and recovery following Hurricane Maria using multitemporal lidar data"

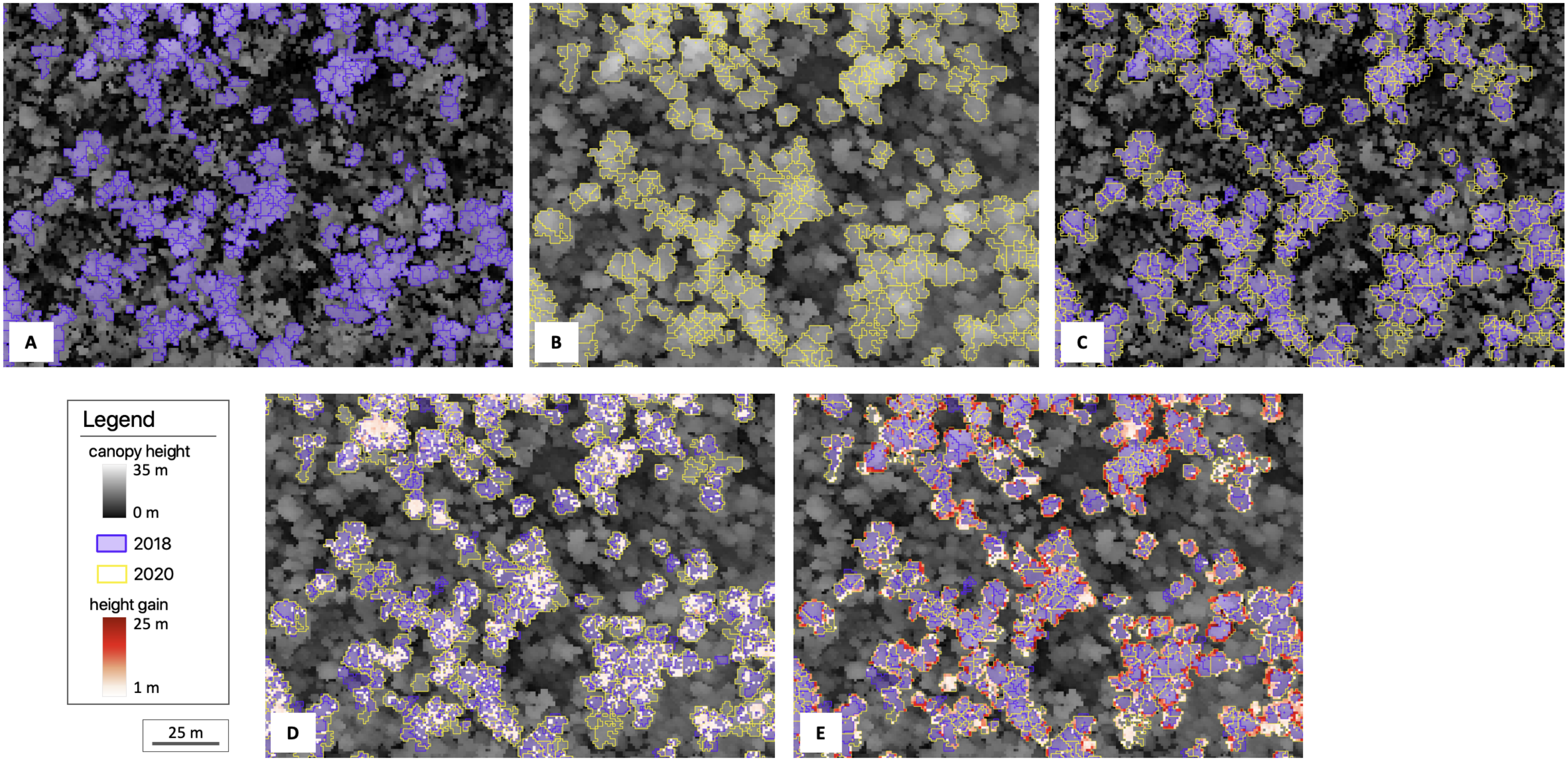


**Figure S1.** An example subset of the MD focus area showing the horizontal extent of surviving tree crown objects (≥19.7 m, the pre-hurricane mean canopy height in MD) identified in (**A**) the 2018 and (**B**) the 2020 canopy height models; (**C**) the union of these two sets of polygons was used to separate 2018-2020 height gain (**D**) within the horizontal extent of 2018 tree crown objects (i.e. *vertical growth*) and (**E**) outside the extent of 2018 but within the extent of 2020 tree crown objects (i.e. *lateral growth*).

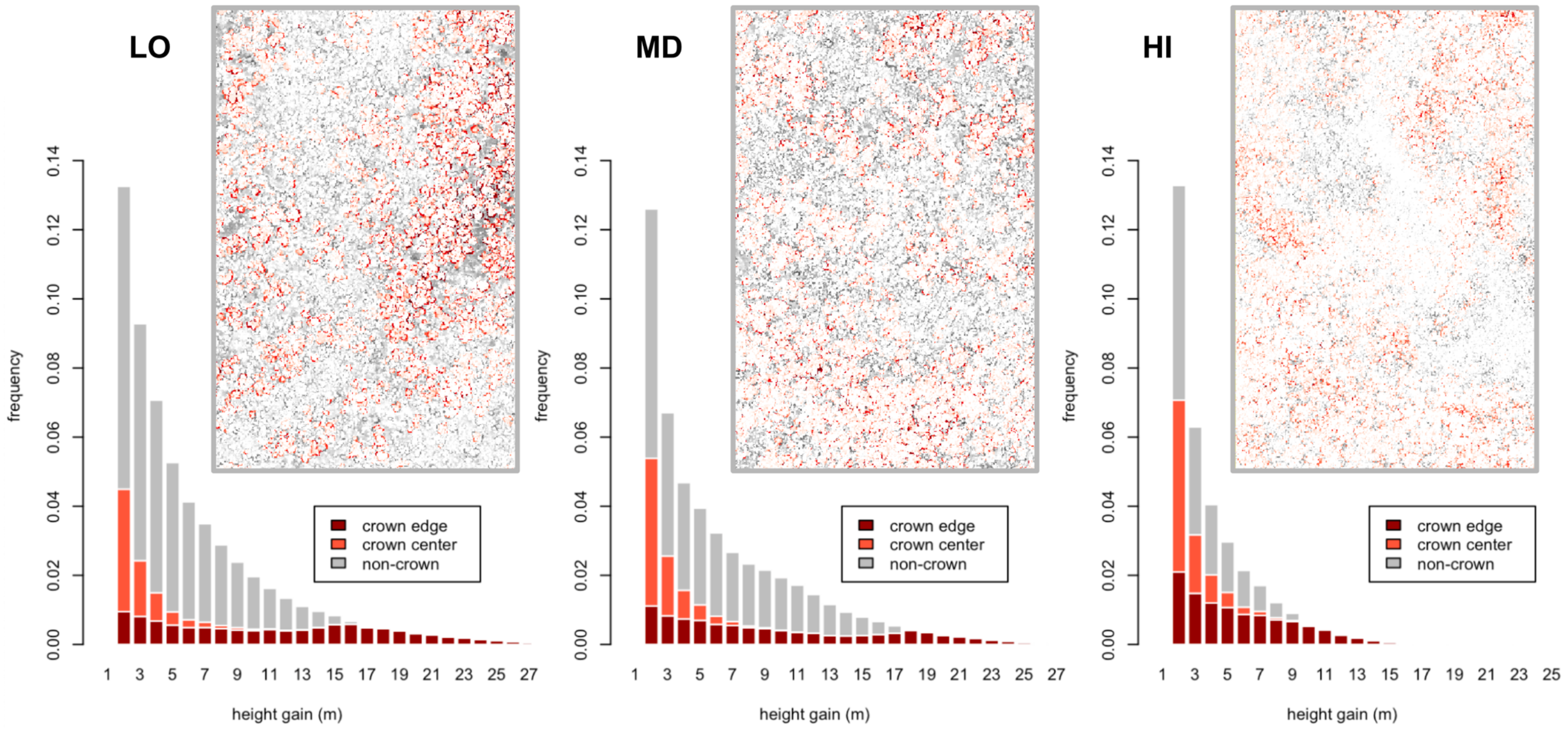


**Figure S2.** Post-hurricane canopy height gains (2018-2020) shown within tree crown objects (center: light red, edge: dark red) versus non-crown areas (gray) in the low-elevation (LO), mid-elevation (MD) and high-elevation (HI) focus areas. Height growth attributed to vertical gains within the horizontal extent of existing crown objects in 2018 (crown centers) was primarily ≤4 m during this interval at all three focus areas (85.1%, 88.3%, and 89.3% for LO, MD, HI, respectively). Estimates of vertical versus horizontal growth for crown edges based on this 4-m height threshold were 22.1% versus 77.9% in LO, 17.7% versus 72.3% in MD, and 44.9% versus 55.1% in HI. Overlapping distributions of height gains in crown centers and crown edges illustrate the difficulty in isolating vertical and horizontal growth patterns, given the complex forest structure following damages from Hurricane Maria.

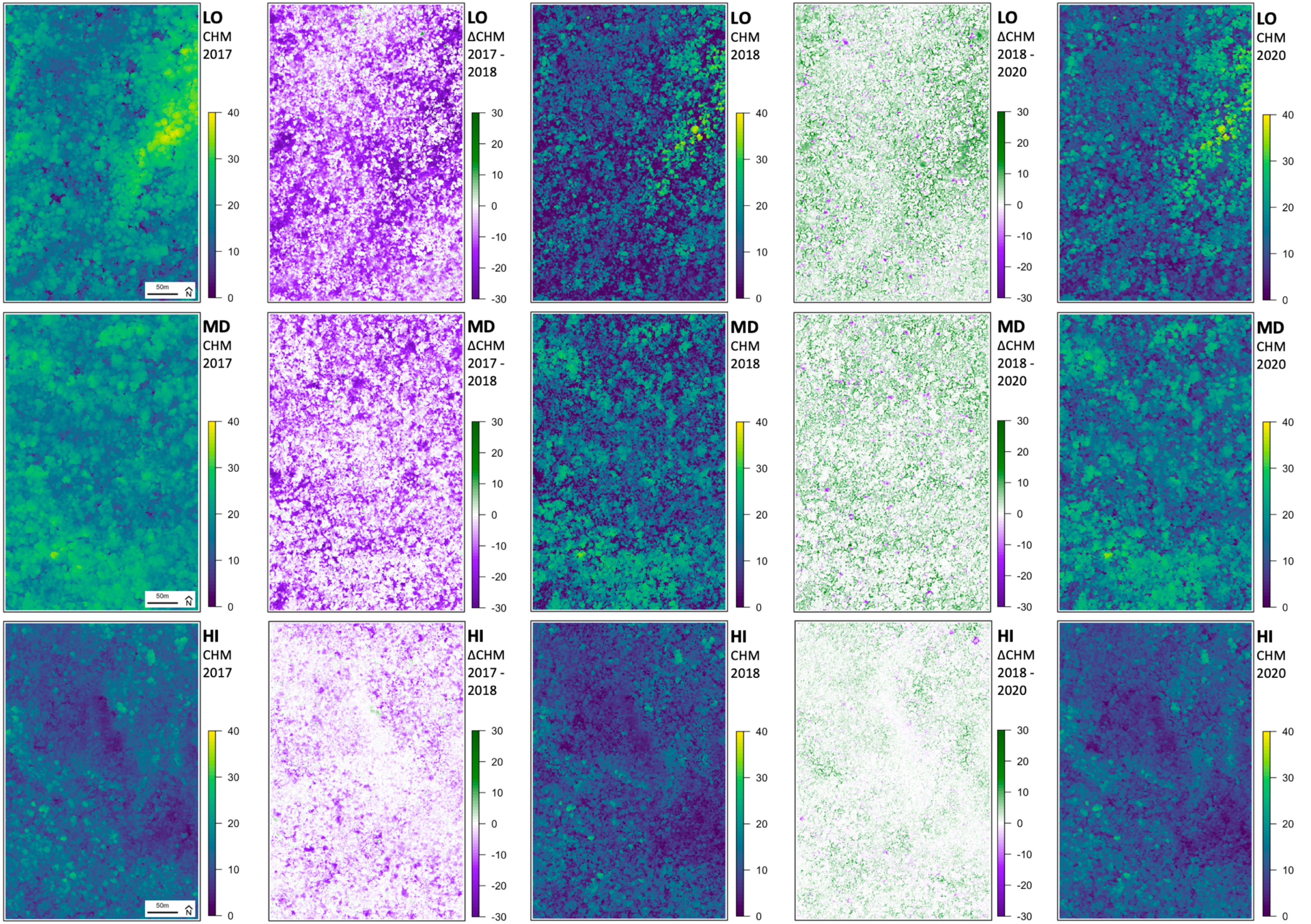


**Figure S3.** Canopy height models (CHM) of the 16-ha focus areas LO (top row), MD (middle row), HI (bottom row) in 2017, 2018, 2020 (1st, 3rd, 5th columns) and respective 2017-18 and 2018-20 change layers (2nd and 4th columns).

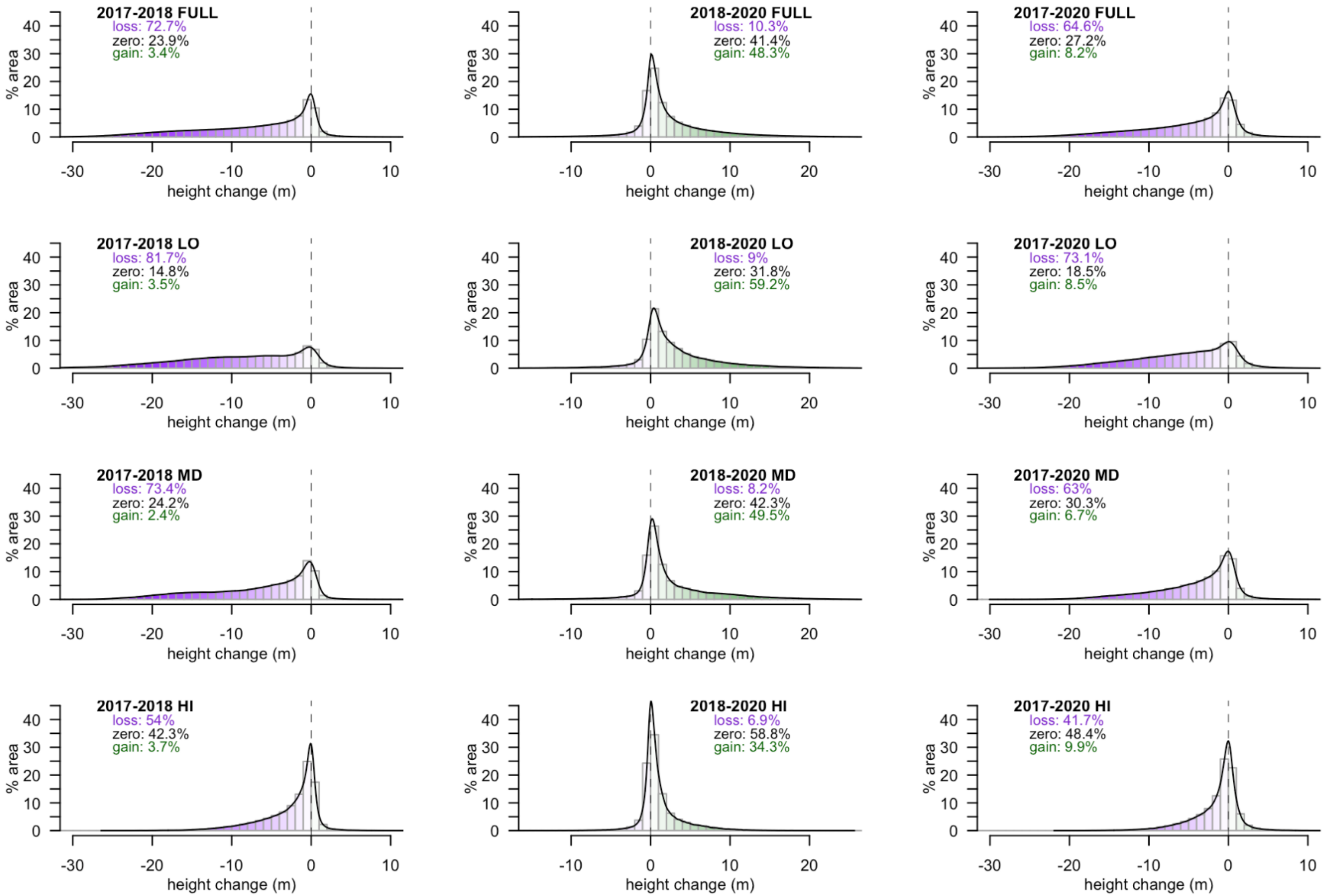


**Figure S4.** Density plots of canopy height changes for 2017-2018 (left), 2018-2020 (middle), and 2017-2020 (right) in the whole study area (FULL, top), and the three 16-ha focus areas at low (LO), mid (BG) and high (HI) elevation (bottom three rows, respectively). The summary statistics in each figure show the percent area of canopy height loss (negative changes of >1 m) in purple, canopy height gain (positive changes of >1 m) in green, and *no change* (where canopy height change was between -1 and +1 m) in gray for each area.
